# In silico evolution of globular protein folds from random sequences

**DOI:** 10.1101/2024.11.10.622830

**Authors:** Harutyun Sahakyan, Sanasar G. Babajanyan, Yuri I. Wolf, Eugene V. Koonin

**Affiliations:** Computational Biology Branch, Division of Intramural Research, National Library of Medicine, National Institutes of Health, Bethesda, MD 20894, USA

## Abstract

The origin and evolution of protein folds are among the most challenging, long-standing problems in biology ^1,2^. Although many plausible scenarios of early protein evolution leading to fold nucleation have been proposed ^3-8^, realistic simulation of this process was not feasible because of the lack of efficient approaches for protein structure prediction, a situation that changed with the advent of powerful tools for fast and robust protein structure prediction, such as AlphaFold ^9,10^ and ESMFold^11^. We developed a computational approach for protein fold evolution simulator (PFES) with atomistic details that provide insights into the mechanisms of evolution of globular folds from random amino acid sequences. PFES introduces random mutations in a population of protein sequences, evaluates the effect of mutations on protein structure, and selects a new set of proteins for further evolution. Repeating this process iteratively allows tracking the evolutionary trajectory of a changing protein fold that evolves under selective pressure for protein fold stability, interaction with other proteins, or other features shaping the fitness landscape. We employed PFES to show how globular protein folds could evolve from random amino acid sequences as monomers or in complexes with other proteins. The simulations reproduce the evolution of many simple folds of natural proteins as well as the evolution of distinct folds not known to exist in nature. We show that evolution of a stable fold from random sequences, on average takes 3 to 8 amino acid replacements per site, suggesting that simple but stable protein folds can evolve relatively easily. These findings could shed light on the enigma of the rapid evolution of protein fold diversity at the earliest stages of life evolution. PFES tracks the complete evolutionary history from simulations that describes intermediate states at the sequence and structure levels and can be used to test versatile hypotheses on protein fold evolution.

## Introduction

The origin of life, one of the most important problems in all of science, is closely connected with the origin and evolution of proteins that are central to all biological processes ^12^. Proteins have the most diverse functions among biological macromolecules and are involved in all cellular processes. Protein function is determined by its structure, or fold, which is formed by folding the linear amino acid chains into a specific three-dimensional shape ^13-15^. Origin and evolution of protein folds remain extremely challenging problems because the methodology required for thorough exploration of evolutionary trajectories is still lacking despite major advances in evolutionary genomics, phylogenetics, bioinformatics, and theoretical biophysics during the last decades. Although many potentially plausible scenarios of protein evolution have been proposed, these are primarily based on simplified, in particular, lattice models that adopt reduced amino acid alphabets and cannot be directly tested experimentally ^16-18^. Nevertheless, many hypotheses converge on the scenario of proteins originating as small peptides with random sequences that gradually evolved into more complex structures with distinct folds ^2-8,19^. A detailed scenario has been proposed in which proteins would evolve in the primordial RNA world when ribozymes using amino acids and short peptides as cofactors gained peptidyl transferase activity, and gradually, this mechanism evolved into the translation system ^20^. An alternative hypothesis, based on a simplified computational model of polar and nonpolar monomers, posits that at the earliest stages of protein evolution, small random peptides formed foldamers, which had the capacity to interact and elongate by self-catalyzing peptide bond formation ^21^.

Until recently, simulation of protein evolution with all-atom models aiming at testing such hypotheses was not feasible because of the lack of fast and accurate approaches for protein structure prediction. The development of machine learning-based tools for fast and robust protein structure prediction including AlphaFold, RoseTTAFold, and ESMfold has changed this situation ^9-11,22^. Taking advantage of these protein structure modeling methods, we developed a toolkit for protein fold evolution simulation (PFES) to simulate and analyze protein fold evolution with atomistic details under different conditions. Using PFES, we show how a protein fold could evolve from random sequences as a monomer or in a complex by interacting with another protein. From multiple protein evolution simulations, we provide an estimate of how many mutations are required for a protein fold to nucleate from a random sequence.

### Protein fold evolution simulation

The main design of PFES is straightforward: i) random (or quasi-random) mutations are introduced into a population of polypeptide sequences, ii) the effect of mutations on protein structure is evaluated, and fitness scores are calculated, and iii) a subset of polypeptides from the given generation is selected for the next iteration (Figure 1A). By iteratively introducing and evaluating thousands of mutations in a population of evolving random amino acid sequences, one can observe how protein structure nucleates and changes through large-scale conformational rearrangements, switching from one fold to another. The simulation starts with a population of size ***N*** that can be created by generating ***N*** random sequences or mutating one random or predefined sequence ***N*** times to create ***N*** mutated variants of the initial sequence. Possible mutations and probabilities for each mutation are provided in several “evolutionary dictionaries” that contain information about allowed mutations and their probabilities (see Methods for details).

**Figure 1.**
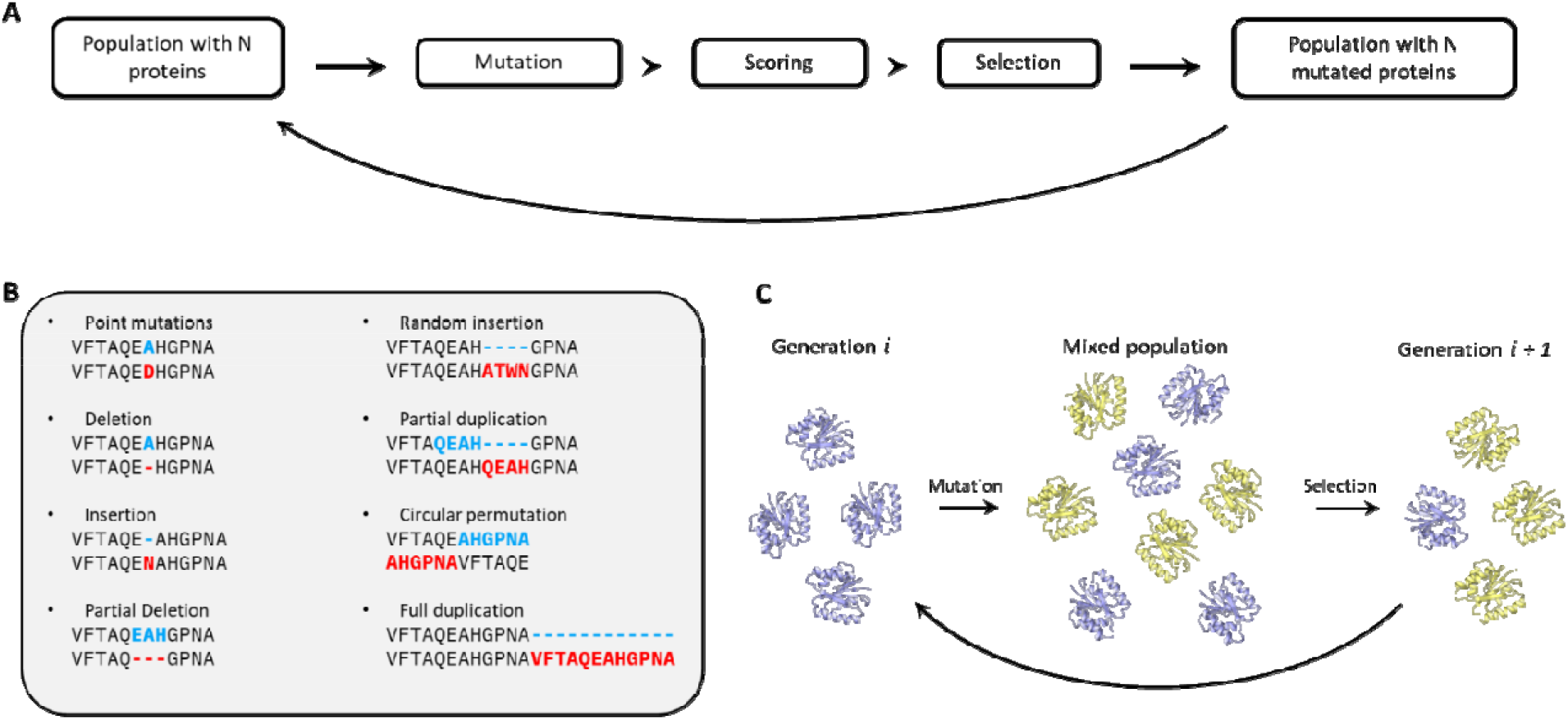
Workflow of protein fold evolution simulation. **A)** Each protein in a population mutates, and the effect of each mutation on protein structure is evaluated to calculate a fitness score; the next generation of proteins is selected based on the fitness scores. **B)** Possible mutation types that occur during the simulation. **C)** Mutation and selection mechanism. All proteins in the population mutate, creating a mixed population of size ***2N*** from which the next generation is selected.

The probability of amino acid substitutions can be uniform with flat rates, that is, any substitution is equally probable. Alternatively, the substitution probabilities can be defined by the codon frequencies or the occurrence of amino acids in natural proteins or in any other manner, for example, using substitution matrices such as PAM or BLOSUM ^23,24^. In addition to the amino acid substitutions, several types of non-point mutations affecting protein length are allowed, including single amino acid insertion or deletion, multiple amino acid deletion, insertion of several random amino acids, partial or complete duplications, and circular permutations (Figure 1B).

The effect of each mutation is evaluated at the next step, based on the structure prediction with ESMfold ^11^. Although any other method can be used in this workflow, ESMfold has a good tradeoff between speed and accuracy, allowing simulation of multiple generations of proteins where changes in the protein fold or other potentially important events can be observed. The final fitness score, calculated for each protein in the population, is a multiplicative function consisting of several terms (see Methods) that reflect the predicted model quality and fold stability, which is one of the key constraints in protein evolution because most proteins require a stable fold to be functional, and moreover, many unfolded proteins are toxic to the cell ^25,26^.

By introducing mutations into all sequences in the population, we create a mixed population of size ***2N*** containing the original and mutated sequences (Figure 1C). For the next generation, ***N*** proteins are selected via weak or strong selection. In the strong selection mode, ***N*** proteins with the highest fitness score are passed from generation ***i*** to ***i+1***, that is, only the fittest half of the mixed population survives to form the population of the next generation. Although strong selection is relatively rarely observed in biological evolution, it has helpful application in the PFES framework for quickly optimizing the already evolved proteins by maximizing their fitness scores. The weak selection mode represents a more plausible evolutionary process with stochastic selection, which allows random drift and makes possible deeper exploration of the fold space. The probability of a protein survival, in this case, is proportional to the fitness score and can be additionally regulated by a β-factor to change the selective pressure (see Methods).

### Evolution of protein folds from random sequences

A series of simulations with varying parameters were run, representing different scenarios in an attempt to understand how protein folds could evolve. In the simplest case, a simulation in the stochastic (weak) selection mode started from 100 random peptide sequences consisting of 24 amino acids each. The initial set of peptides emerging from 100 different mutations of a random sequence forms the first generation. In the simulation, sequences randomly mutate, and under selection, favorable mutations gradually accumulate, determining the protein structure. Initially, favorable mutations result in the formation of contacts within the evolving peptide, and later, when enough contacts are established, the peptide adopts a distinct fold with defined secondary structures. Typically, this fold is very simple, such as alpha/beta-hairpin, helix-turn-helix (HTH), or WW domain, as illustrated in Figure 2A. As a rule, the nucleated structure had a low score, indicative of instability of the fold, but as the population evolves, more stable folds originate with higher pLDDT and pTM scores. Once a minimal stable structure is fixed, it goes through small changes where, for hundreds and thousands of mutations, the fold architecture does not undergo major modifications although the sequence can drift and change substantially. This process can be described as the evolving protein reaching a local peak on the fitness landscape ^27^ where only small and gradual changes in the loops or minor modifications of the secondary structure take place, usually leading to more compact and stable structures. More noticeable changes in the structure are typically associated with small insertions, which also occur mainly in the loops. For example, the C-terminal loop in one of the evolving proteins started forming a small 3_10_ helix that grows and transforms into a short alpha-helix (Figure 2A,B; Movie S1).

**Figure 2.**
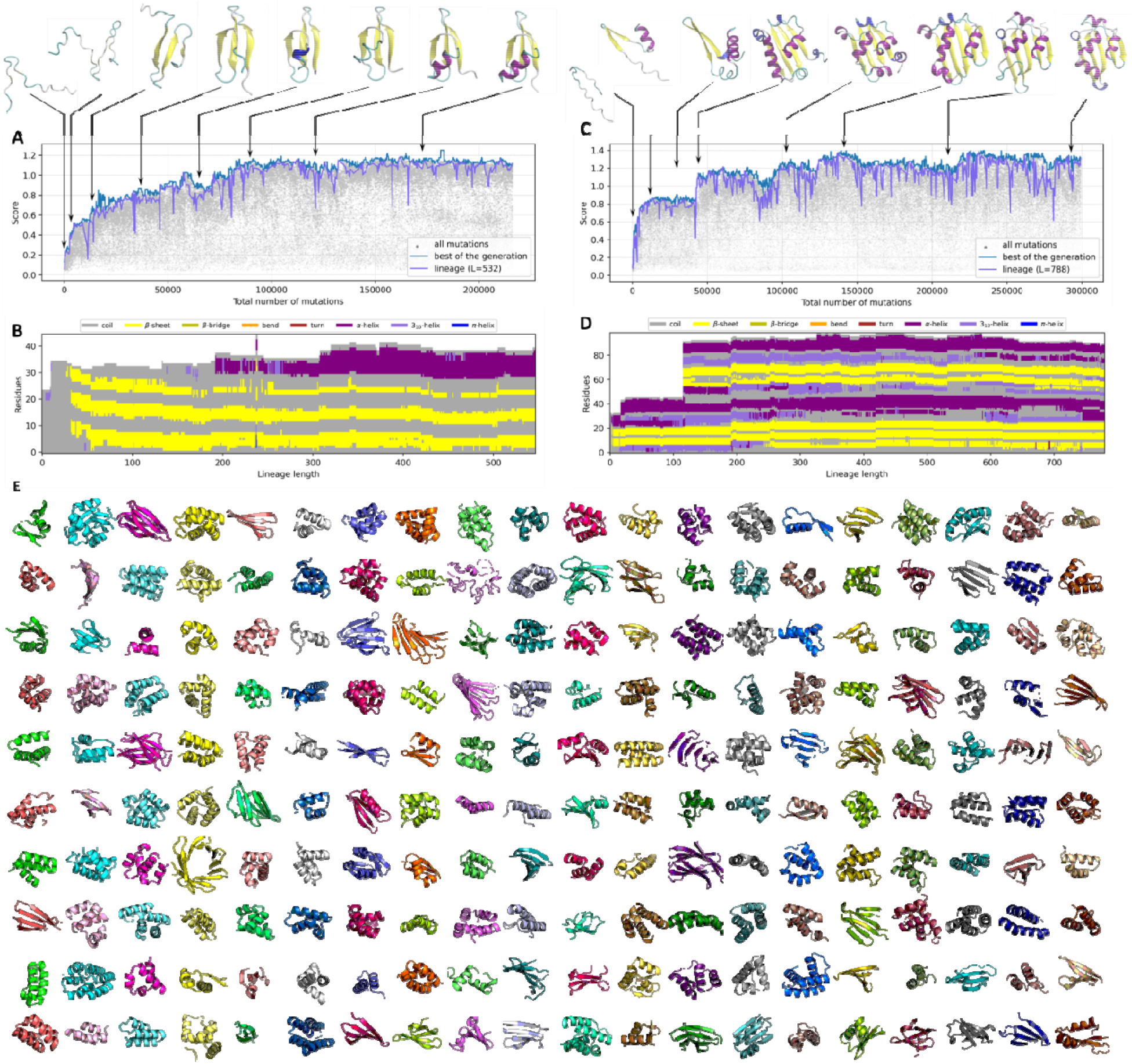
Evolution of protein folds in PFES. **(A)** The fitness score plotted for all sequences (grey), the best sequences in each generation (blue), the lineage of the best sequences from the last generation (violet), and intermediate structures of a gradually evolving protein representing different points of the evolutionary trajectory. **(B)** Changes in the secondary structures after each mutation in the lineage. **(C)** Fitness score and representative structures of a protein evolving via complete domain duplication. **(D)** Corresponding changes in the secondary structures. **(E)** Protein structures evolved in the simulations. The models are arranged in the order of simulations.

Each simulation yields a unique evolutionary trajectory, leading to a different protein in each case. The simulations described above were repeated 200 times, and mostly, the evolved protein had a distinct fold, although some generic similarities were observed (Figure 2E). Clustering of the final folds based on a minimal TM-score of 0.5 showed that the majority of folds (110 of 200) were unique, whereas the remaining ones formed clusters, of which the largest consisted of proteins with simple folds, such as HTH, beta-sandwiches, beta-sheets, and alpha/beta-hairpins. Clustering these proteins at 30% sequence identity yielded only singletons, indicating that the evolving proteins shared virtually no sequence similarity. PSI-BLAST search in the NCBI NR database yielded no significant sequence matches among natural proteins except for the protein from simulation #29, which showed significant similarity to some uncharacterized Zn-finger proteins. Protein folds consisting mostly of alpha helixes were the most abundant in our simulations (Figure S1A), which can be explained by the mechanism of their evolution. Once a small alpha helix is nucleated, it continues to grow gradually with insertions at the termini or in the middle of the helix until a critical length is reached. Duplications often lead to the formation of more complex structures, such as an alpha-hairpin from a single helix or an alpha-helical bundle from an alpha-hairpin. In evolutionary simulations, simple beta folds also reoccurred multiple times, and the evolution mechanism of these folds often showed a distinct pattern. A beta-hairpin evolved from random sequences duplicated fully or partially several times, forming a larger beta-sheet, which later folded into a beta-sandwich or jelly-roll-like structure. In some cases, this process leads to the evolution of beta-barrel-like structures. For example, the proteins from simulations #188 and #197 produced the SH3 fold, an ancient domain that is widespread in all life forms, particularly among translation system components ^28^.

Structural search with Foldseek ^29^ in PDB, AlphaFold/UniProt50, and MGnify-ESM30 databases yielded multiple hits with 0.8 or greater coverage of the queries and mainly partial coverage of targets. This structural search showed that 82 of the 200 structures evolved in the simulations had structural analogs among natural proteins, with a Foldseek probability of at least 0.95, including 23 hits from PDB (Table S1). The largest number of hits involved simple beta-sheets and other proteins with the mostly-beta folds. For instance, the search with the SH3 domain from simulation #188 recovered 2001 predicted and experimentally solved structures closely resembling this fold. Interestingly, a structurally similar but topologically distinct protein evolved in simulation #197 yielded only 92 hits, although it has a TM-score similarity of 0.68 with protein #188. Other examples with a more complex fold include the protein from simulation #194 resembling low-density lipoprotein receptor chaperone boca (pdbid: 3ofeB) ^30^, with TM-score 0.75 and RMSD 1.9 Å. Protein evolved in simulation #18 showed similarity with a fragment of V-ATPase subunit C (pdbid: 4dl0) with a TM-score of 0.73 and RMSD of 1.99 Å, although the respective sequences shared only 7% identity. Notably, several small proteins with simple folds, such as protein from simulation #100, which resembled a fragment of CRISPR-associated protein Cas5, in addition to the high structural similarity, also showed a relatively high sequence similarity (35-40%) although not detected by sequence search (Figure S3). The complete list of Foldseek hits is provided in Table S2.

Many details of protein evolution can be directly extracted from the simulations for further analysis, including a complete phylogenetic tree containing information about each mutation and all ancestral sequences. The examples in Figures 2A and 2C show the highest score for each generation and the full lineage of the protein with the highest score from the last generation. Although, at some points, this lineage overlapped with the best score of an intermediate generation, the trajectories of the ultimate winner and that of the best intermediate scores were different. Given that weak selection is a stochastic process in which a high fitness score increases the probability of survival but does not guarantee it, the protein with the highest score does not necessarily pass to the next generation. Conversely, variants with better scores often could not be found for several generations, and the fittest variant from the previous generation was repeatedly selected. As a result, the number of unique sequences in the lineage is always smaller than the number of generations. Detailed information about the changes in the protein sequence and the corresponding structural changes can be readily extracted from the lineage leading from the initial random sequence to the final sequence of the last generation with the best score as can be visualized at the level of secondary (Figure 2B, D) or tertiary structure (Movie S1, S2).

Although sequence similarity searches provided limited or no information for the artificially evolved proteins, for 71 of the 200 evolved proteins, AlphaFold2 nevertheless produced structural models with mean pLDDT > 80. The quality of the AlphaFold predictions strongly depends on the number of proteins homologous to the query that can be identified by sequence similarity search. Therefore, proteins with novel, unique folds from our simulation often could not be modeled with AlphaFold2, even when ESMfold predicted these structures with high confidence. For instance, the protein evolved in simulation #2 has a novel fold, which was not observed among the natural proteins. We ran a short simulation with strong selection to optimize this fold, which resulted in 72 mutations substituting 47% of residues in the initial sequence and improving the mean pLDDT from 83.5 to 91.1. The same optimized sequence predicted with AF2 had a mean pLDDT of only 44.7 (Figure 3A). However, when evolutionary information extracted from the simulations was provided as a multiple sequence alignment, AF2 dramatically improved performance predicting a structure for the same sequence with a mean pLDDT of 95.9, that is, a confidence score even higher than that obtained initially with ESMfold (Figure 3A). Protein from simulation #19 is another example with a novel fold, which after optimization had a mean pLDDT of 94.3 by ESMfold, whereas AF2 gave a pLDDT score of only 54.6. However, when information on residue conservation and coevolution from PFES was provided, AF2 predicted the same structure as ESMfold but with a dramatically improved mean pLDDT of 95.5.

**Figure 3.**
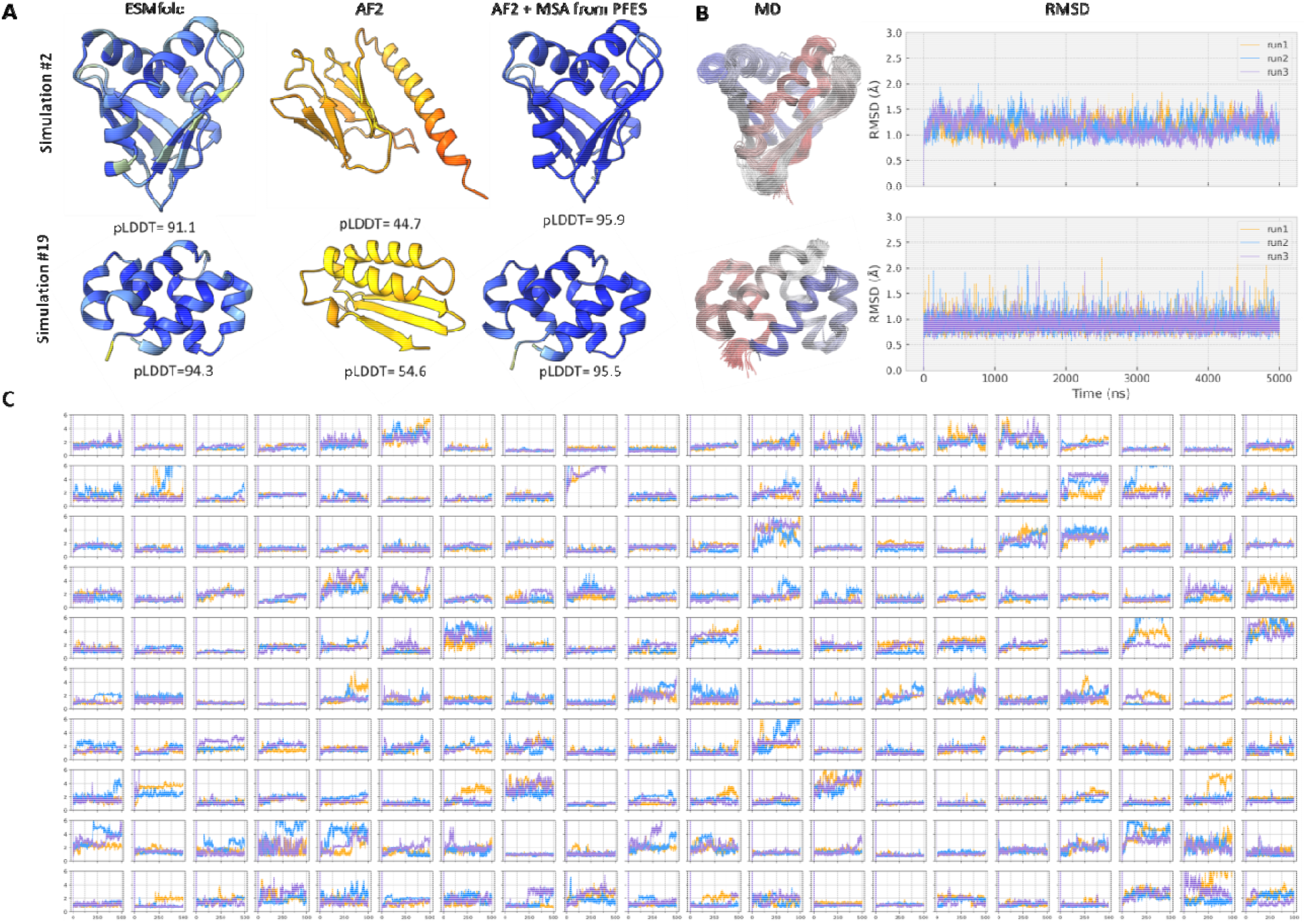
Stability of the evolved proteins. **(A)** Structures of proteins from simulations #2 and #19, predicted with ESMfold, AF2, and AF2 using an MSA extracted from PFES. **(B)** Superimposed snapshots from MD simulations colored from C (red) to N (blue) termini and their RMSD fluctuations over time, showing the stability of these proteins during the MD simulations. **(C)** Backbone RMSD fluctuations for all proteins measured after MD simulations. Plots are ordered in the same way as proteins in Figure 2E.

More dramatic changes in the evolving protein structures often resulted from partial or complete sequence duplications, one of the primary modes of gene evolution ^31,32^. Although duplications are rarely fixed, they often lead to a major increase in fold complexity because after duplication, each of the two copies evolves separately. After the duplication, once the new fold was established, changes in the structure were mostly incremental, affecting the loops or small secondary structure elements. Such a duplication event is well illustrated by the simulation presented in Figure 2C, where a full duplication occurred after 400 generations, shaping a completely new fold with a beta-sheet and several alpha-helices. After duplication, the protein fold did not change its architecture during the rest of the simulation although there were minor changes in the loops that transformed into small alpha-helices, and an additional beta-strain was formed (Figure 2D, Movie S2).

Duplications seem to be essential for the growth of evolving proteins, given that, in a broad variety of organisms, the rate of deletions is higher than the rate of insertions ^33-35^. Under this deletion bias, proteins would shrink with time, contradicting the fact that, at least, in many evolutionary lineages, protein complexity generally increases in the course of evolution. Furthermore, an increase in complexity should have been essential at the earliest stages of protein evolution. In our simulations, proteins were increasing in size most of the time due to selection, even when deletions were more frequent than insertions (Figure S2). Apparently, a lower insertion rate is compensated by the selection for the fitness score, which tends to be higher for larger proteins, and duplications provide an opportunity for a fast increase in fold complexity that is fixed if the duplicated structure is stable enough.

Despite high confidence scores obtained with ESMfold and AlphaFold2, it remained unclear how stable the proteins evolved in simulations were, especially considering the non-natural origin of MSAs that were used to improve AF2 predictions. To address this question, we performed a series of all-atom molecular dynamics (MD) simulations for all artificially evolved proteins with 3 repeats in an explicit water environment for a more robust estimation of protein stability using physics-based simulations that do not depend on evolutionary or any other information except the predicted protein atom coordinates (see Methods). We found that 163 of the 200 evolved proteins remained stable in all three MD simulations with a backbone RMSD below 2.5 Å and a standard deviation of 1, even though 101 of these proteins had unique folds not detected among natural proteins (Figure 3B, C). The absence of dramatic deviations for the majority of the predicted structures during the simulation provides additional confidence that the predicted structures are stable.

### Number of mutations required for fold nucleation

One crucial question that can be addressed with PFES is how many mutations are needed to change a random peptide into a protein with a stable fold. Some small proteins have very simple structures, such as a beta-hairpin or an alpha-helix, but the evolution of larger proteins, usually containing more than 50 residues, strongly depends on the rates of duplication and insertion that are necessary to elongate the protein when the simulation starts from a small peptide. To circumvent the parameter search problem, we started simulations from 50 amino acids long polypeptides allowing only point mutations and single amino acid indels to maintain approximately the same length throughout the simulation. In this regime, we ran additional 200 independent simulations with a fixed population size of 100 members as in the previous runs until one of the peptides in the evolving population became stable, and as a condition for protein stability, pLDDT of 0.85 and pTM of 0.75 was selected. When the simulations reached the stopping conditions, we extracted lineages from each simulation to calculate the number of mutations necessary to evolve a stable protein from a random sequence. In this case, most of the evolved proteins had all-alpha folds such as HTH or helical bundles, and proteins with mixed folds containing both alpha helices and beta sheets, evolved more often than all-beta folds (Figure S1A). For most random sequences, it took on average 145 mutations to evolve into a stable fold, that is, ∼2.9 mutations per site (Figure 4A), considering that the length of the sequences did not change much during the simulations (Figure S1B). However, depending on the initial sequence and the evolution trajectory, the number of necessary mutations could vary dramatically. Whereas in some cases, only 10 mutations would suffice, in others, more than 500 mutations were needed to evolve a stable fold. It should be noted that this is the number of mutations only in the lineages that represented the direct path from the initial random sequence to the final protein. Many more mutations are needed to find this path: on average, it took 531 generations (or 53,100 mutations in total) to evolve a stable fold from a random sequence (Figure 4B), that is, only 0.003% of the mutations were fixed.

**Figure 4.**
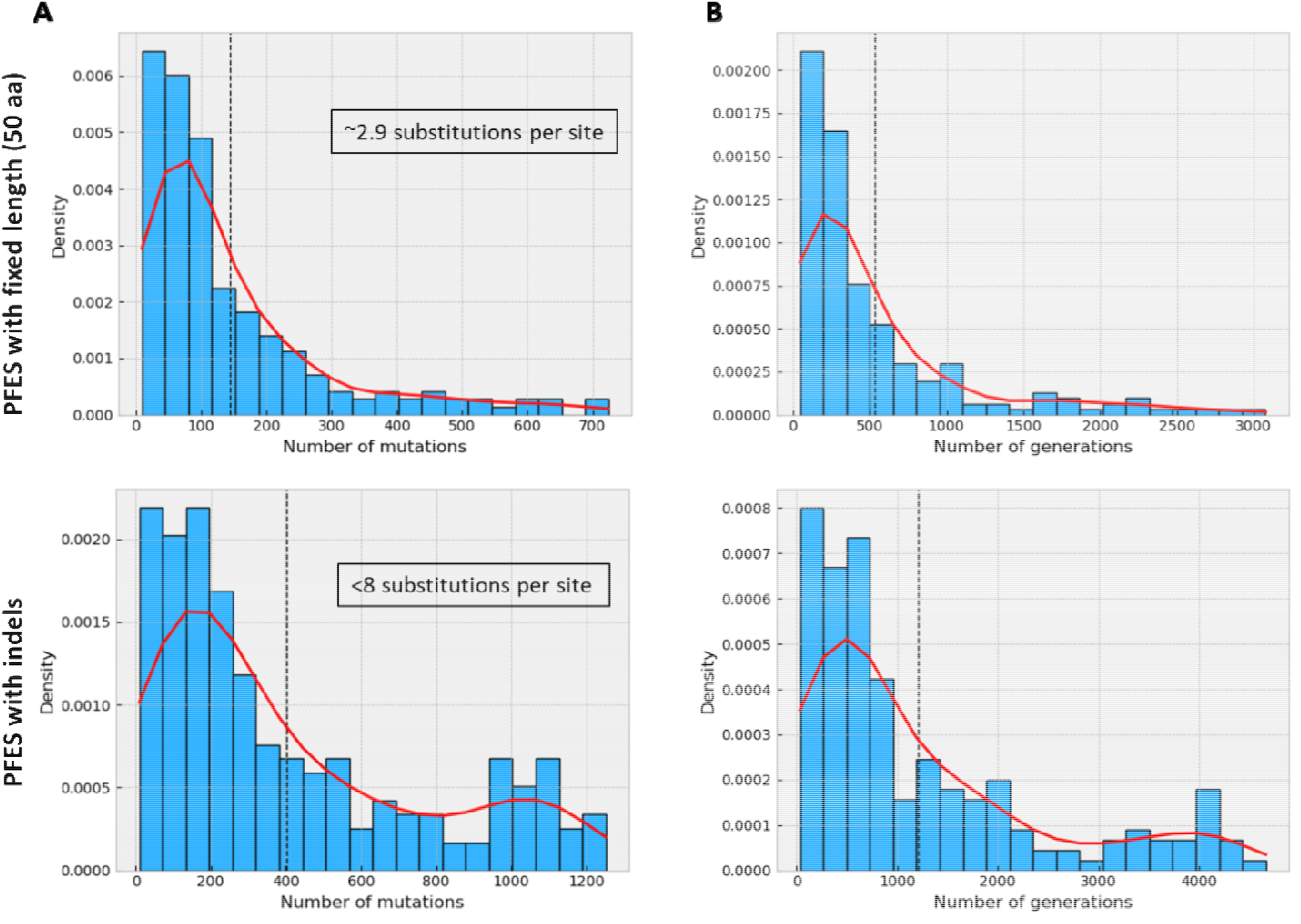
Number of mutations to fold nucleation estimated from simulations with fixed chain length or with indels. **(A)** Distribution of the number of mutations required for the transition of a random sequence into a stable fold. **(B)** Number of generations to fold nucleation. The dashed line shows the mean of the distribution, the red line shows kernel-density estimate using Gaussian kernels.

We also calculated how many mutations or how many generations it took for an evolving protein to nucleate into a stable fold from the previously described simulations with indels. In these simulations, most proteins adopted all-alpha folds (Figure S1A), and it took, on average, ∼400 mutations or ∼1240 generations to evolve a stable protein from a pool of random 24-residue peptides (Figure 4A, B). The average size of the final proteins was ∼56 residues (Figure S1B), but because the proteins were smaller at the beginning of the simulation and grew gradually via insertions and duplications, there were less than 8.1 mutations per site.

These simplified experiments show how, using simulations, we can estimate the number of mutations to protein fold nucleation. Many other parameters, such as population size, selection strength, mutation types, and their probabilities, will evidently affect this estimation. Moreover, the protein stability criteria that we used (pLDDT of 0.85 and pTM of 0.75) are also arbitrary, and using other, stricter, or softer conditions will also change these estimates.

### Evolution of proteins with interacting partners

Although protein fold stability is a necessary condition for most proteins to be functional, proteins do not exist in isolation but typically form a complex network of protein-protein interactions. Some of such interactions fulfill critical functions, which creates extra evolutionary pressure, in addition to the requirement of protein fold stability. To reach high fitness, a protein should often not only be stable but must also interact with specific, functionally important partners. We simulated a process whereby an evolving protein interacted with another protein that did not change through the simulation. The following example illustrates how random sequence evolves by interacting with a small DNA-binding domain with the HTH fold (bacterial cell cycle regulator GcrA) ^36^. The simulation of 3000 generations was performed with stochastic selection, a population size of 100, and initial random sequences of 24 residues in length. The stable protein fold evolved during the first 250 generations and did not dramatically change thereafter (Figure 5A). While evolving, the protein interacted with different parts of the HTH domain, and only after the 2000th generation did the interaction stabilize as more contacts formed between the two chains (Figure 5C). The C-terminal helix of the HTH carrying a positive charge was essential for the interaction with the negatively charged major grove of the DNA, and the evolved protein that contained a negatively charged pocket mimicked DNA (Figure 5A), binding the HTH domain via electrostatic interactions.

**Figure 5.**
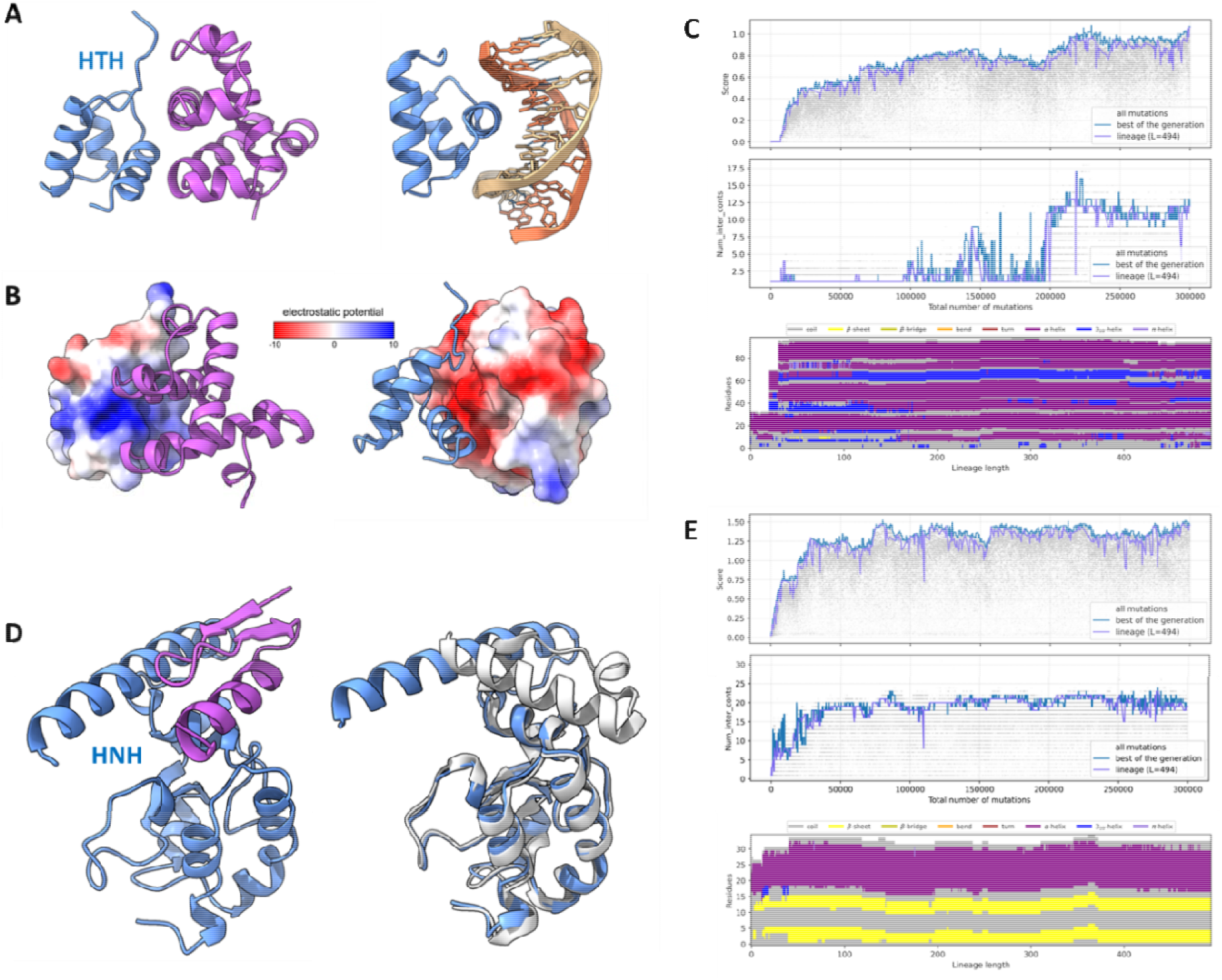
Evolution of a protein fold in complex with a binding partner. **(A)** The structure of a protein evolved from a random sequence (magenta) binding HTH of GcrA and the same HTH domain in a complex with dsDNA (PDBID: 5YIW). **(B)** Electrostatic surfaces of the HTH charged positive (left), and the protein evolved in the simulation charged negative (right). **(C)** Fitness score and the number of contacts between two chains plotted for all sequences (grey), the best sequences in each generation (blue), and the lineage of the best sequences from the last generation (violet). The effect of each mutation on the secondary structures is plotted for all mutations in the lineage. **(D)** The structure of an evolved protein in complex with HNH domain (blue). On the left is the conformation of the HNH domain in complex with the evolved protein (blue), superimposed with a free HNH domain that does not interact with other proteins (PDBID: 6O56, grey). The evolved protein is hidden for clarity **(E)**. Changes in the score, the number of contacts between two chains, and the secondary structures as described above.

We also ran a simulation with the same parameters as in the example above to reproduce the evolution of an anti-CRISPR protein (Acr) from a random sequence. Most of the numerous identified Acrs are small, fast-evolving proteins that interfere with different components of the bacterial and archaeal CRISPR-Cas adaptive immune system ^37,38^. In this simulation, the protein was evolved from random sequences by interacting with the conserved HNH nuclease domain of the Cas9 protein that is essential for the target DNA cleavage and is targeted by several natural Acrs ^39,40^. The random peptide evolved into a simple zinc finger fold with two beta-strands and an alpha-helix (beta-beta-alpha zinc finger, BBA) binding to a hydrophobic region of HNH consisting of residues 775-792, 802-818, and 887-893 (Figure 5D). As in the previous cases, the major changes in the fold of the evolving protein occurred in the beginning of the simulation, and after ∼500 generations, the fold and the interaction mode between the two proteins changed only marginally (Figure 5E). The evolved protein was not charged as in the previous case, and the interaction between the two proteins was stabilized by the hydrophobic effect. The evolved protein bound the alpha-helix formed by residues 775-792 and changed its conformation upon binding (Figure 5B). The binding of the evolved BBA protein did not directly obstruct the access to the catalytic site, as it does, for instance, in the case of AcrIIC1 ^41^, and did not block the PAM-recognition site like the DNA mimicking AcrIIC2, AcrIIA4, and AcrIC5 ^42^ but nevertheless, this interaction could lead to incorrect domain packing in Cas9, affecting its activity.

## Discussion

The emergence of stable, globular protein folds from random sequences is arguably the principal problem of protein evolution and one of the major challenges in the study of the origin of life. Once diverse globular folds evolved, the subsequent evolution of proteins throughout the 4 billion year history of life on Earth was a relatively straightforward and fairly well understood process. Successful attempts to computationally imitate protein evolution have been made previously, but these studies either focused on a particular part of the protein with amino acid substitutions that did not change the protein fold ^43-45^ or were limited to the exploration of simplified models in lattice space and reduced amino acid alphabets ^18,21^. Here, in contrast, we simulated protein evolution with all-atom models, allowing us to observe how protein folds nucleated from random sequences and how the accumulation of amino acid substitutions led to large-scale conformational rearrangements, changing the general architecture of the fold. This critical difference that was made possible by the advent of powerful tools for protein structure prediction, such as AlphaFold and ESMfold, creates the opportunity to recapitulate protein fold evolution in detail not accessible previously.

The principal result of this work is that emergence of simple, stable, globular protein folds from random amino acid sequences is relatively easy. Especially, under strong selection, many simulations readily produce such folds, some of which are closely similar to folds found in natural protein domains, such as HTH, beta hairpins, WW or SH3. On average, we observed that nucleation of a stable fold required about 3-8 amino acid substitutions per site of the evolving amino acid sequence, depending on the size of the initial sequences and the evolutionary regime. This is a substantial but not unrealistic amount of evolution. In different evolutionary lineages, the characteristic rates of protein evolution range between 0.2 and 1.5 amino acid substitutions per site per billion years ^46^. The time available for the emergence of protein folds at the early stage of life evolution was several hundred million years at best, under the latest robust estimates which suggest that well defined protein folds already existed earlier than 4 billion years ago ^47^. Furthermore, there are indications that the emergence of a diverse repertoire of protein folds antedated the formation of the full-fledged translation system, that is, apparently occurred at a stage of evolution directly following the primordial RNA World ^48,49^. However, the above amino acid substitution estimates apply to proteins that evolved under strong constraints imposed on established, optimized folds. It appears highly likely that evolution from random sequences at the primordial stages was much faster. Furthermore, in some of our ‘luckier’ simulations, the change required to reach a stable, globular conformation was far less extensive. Thus, our results suggest that evolution of globular protein folds from random sequences could be straightforward, requiring no unknown evolutionary processes, and in part, solve the enigma of rapid emergence of protein folds. Furthermore, the appearance, in many of the PFES runs, of simple folds closely similar to those found in natural proteins implies that evolutionary trajectories in the folding space are strongly constrained.

The major limitation of PFES is the limited accuracy of protein structure prediction methods. Although ESMfold provides a good tradeoff between speed and accuracy, it is questionable how biophysically realistic the predicted structures are, especially, structures with low confidence scores that represent the intermediate states of protein fold evolution. Nevertheless, proteins in the native environments are not fixed constructs as experimental methods capture them but rather exist in a folding-unfolding equilibrium ^50^. The fraction of the unfolded protein is determined by the thermodynamic stability of the protein, that is, the folding free energy ^51^. The same fold predicted with ESMfold for different sequences can have different confidence scores, which might reflect the fraction of folded vs unfolded proteins. Thus, a protein with a low confidence score is likely to maintain that fold transiently, whereas most of the time, such a protein is unfolded. If so, such sequences that possess only minimal functionality nevertheless could be subject to selection that will gradually stabilize the proteins, shifting the equilibrium towards a greater fraction of folded, stable proteins, with higher ESMfold scores. This is indeed the pattern of de novo protein evolution that we observed in PFES simulations.

Due to computational power limitations, we applied PFES to explore only a limited region of the protein evolution parameter space in a relatively small number of runs. However, PFES is a rich, flexible framework that, in principle, provides for a deep and broad computational study of protein evolution. With further development of PFES combined with improved prediction of protein structure and interactions with other proteins as well as nucleic acids, more realistic reproduction of primordial events at the origin of life will likely become feasible.

## Methods

### Protein fold evolution simulation

Protein fold evolution simulation (PFES) imitates the process of protein evolution by mutating a population of proteins, evaluating those mutations, and selecting the next generation of further evolution (Figure 1). There are 3 predefined rates for mutation that can be additionally adjusted in an arbitrary way. In the presented simulations, the rate of single amino acid substitutions was uniformly distributed, and the probabilities used for non-single mutations are provided in Table S3. The size of the population (***N***) is defined at the beginning of the simulation and remains fixed. Each protein in the simulation mutates once, and the next generation is selected from the mixed population of original and mutated proteins of ***2N*** size. Strong or weak (stochastic) selection can be used in the simulations to determine which proteins will be transferred to the next generation. Strong selection (Eq 1) mode is deterministic, and only the best half of the mixed population forms the next generation.

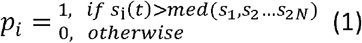

where *s*_*i*_ is fitness score of sequence i. Note that here, a given sequence may be selected at most once. However, the same sequence can appear in the population due to reoccurring mutations.

The probability in the stochastic selection used in these simulations was calculated assuming that *p*_*i*_ is defined by relative fitness advantage:

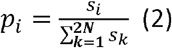

Where 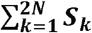 is the sum of all scores. In this case, the same sequence can be selected multiple times depending on its score. The selection strength in the stochastic selection mode can be additionally regulated by a β factor as in the Gibbs distribution:

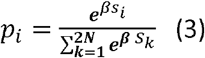

With *β* = [0, ∞], where *β* = 0 mean no selection that is 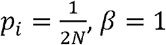, resembles stochastic selection as in Eq (2), and *β* > 5, starts behaving as strong deterministic selection as is Eq (1) (Figure S4).

The effect of each mutation is evaluated based on the structure predicted by *esmfold_v1* pretrained model with one recycle ^11^. This is the bottleneck of the algorithm, and structures from each generation are split into several batches containing multiple sequences, which are predicted together for optimal usage of the GPU. While the next batch is being predicted, the algorithm calculates scores for the previous batch using a CPU, thus running two processes in parallel to optimize the performance. The final fitness score is a multiplicative function that includes pLDDT, pTM, contact density (CD), and constraints if they are applied Eq 4.

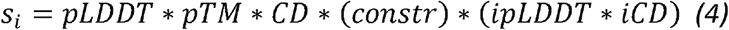

The first two terms (*pLDDT, pTM*), obtained directly from the ESMfold prediction, show the general prediction quality. Contact density (*CD*) is calculated as the number of contacts normalized by the protein length l (4.1). Contacts are calculated between C-beta atoms if Euclidian distance those are closer than 6 Å, residues are located 5 positions apart in the sequence (d1 and d2 in Eq 4.1) and pLDDT for both residues is more than 50, so contacts in the alpha-helices or low pLDDT regions are not counted.

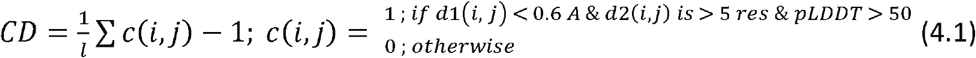

For dimers, the score includes two additional components: *ipLDDT*, which is *pLDDT* on the interface of interacting proteins, that includes residues between chains where C-beta atoms are closer than 6 Å, and *iCD*, which is the number of contacts between chains calculated in the same way and normalized by the length of the evolving protein.

Additional correction in the score can be introduced through constraints for protein or secondary structure length (P_L_ and P_α_, P_β_), which are functions with values from 0 to 1 depending on protein or secondary structure lengths (Eq 5). For instance, if protein length is within the allowed range, the score is multiplied by 1 and does not change, but if it becomes equal to the Lo, the score is multiplied by 0.5.

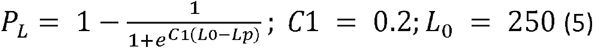

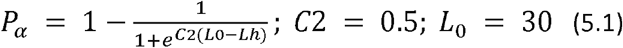

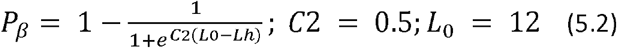

Secondary structure elements are identified from the predicted structures using PSIQUE (P^52^. In the simulation in this work, the secondary structure constraints Lo for beta-sheets was 12 and for alpha-helices 30.

Two sets of simulations were run with 200 repeats in each. The first set included simulations included all possible mutations mentioned above was performed for 4000 generations with stochastic selection and 1000 generations with strong selection at the end. The populations size was 100 and the initial population was started from a random sequence with a length of 24 amino acids. The second set of simulations was performed with constrained length using only point mutations and single amino acid indels. The initial population was started from a random peptide with a length of 50 amino acids, the population size was 100. These simulations were run with stochastic selection until one of the peptides in the generation reached pLDDT of 85 and pTM of 0.75.

PFES is available at github.com/sahakyanhk/PFES

#### Structure similarity search and clustering

GTalign was used for clustering structures with a TM-score of 0.5 and coverage of 0.7 with speed mode 0 ^53^. MMseqs2 with a sequence identity of 30% and bilateral coverage of 0.7 was used for sequence clustering ^54^. Foldseek was used for structure-based search in AFDB/UniProt50, MGnify-ESM30, and PDB with alignment type 3Di+AA and a sensitivity of 9.5 ^29^. Hits were filtered with probability > 0.95 and query coverage 0.8. PSI-BLAST with word size 2 was used for search in the NR database ^55^.

#### Molecular Dynamics Simulations

Molecular Dynamics (MD) simulations were performed using Amber20 software and ff19SB protein forcefield ^56,57^. Proteins were solvated with the TIP3P water model and K^+^/Cl^-^ ions in truncated octahedral boxes with 10 Å buffer between protein and box. The systems were minimized and equilibrated gradually, releasing restraints, and the final simulations were performed at a constant temperature of 298.15 K and pressure of 1 bar using the Langevin thermostat with a collision frequency of 2ps^-1^ and Monte Carlo barostat, respectively. PME approach with a short nonbonded cutoff of 10 Å was used for treating long-range electrostatics. For the final simulations, hydrogen mass repartitioning (HMR) and 4 fs integration timestep was used. MD simulations lasting 500 ns were repeated 3 times for each protein, except for proteins from simulations #2 and #19 which simulations were performed for 5000 ns. RMSD for each trajectory was calculated for backbone atoms with *cpptraj* using the equilibrated structure as a reference ^58^.

### Protein structure prediction

Structures of artificially evolved proteins were predicted using AlphaFold2 v2.3.2 with default parameters ^10^. ColabFold v1.5.5 was used to predict structures with custom MSAs, which were extracted from PFES ^59^.

### Molecular graphics

For protein structure visualization, ChimeraX, Pymol, and VMD were used ^60,61^ https://legacy.ccp4.ac.uk/newsletters/newsletter40/11_pymol.pdf). PFES lineage trajectories showing changes in the protein structure can be visualized in VMD ^61^.

## Data availability

Simulation files are available at https://doi.org/10.5281/zenodo.14061035.

## Code availability

PFES code is available at github.com/sahakyanhk/PFES

## Author contributions

H.S. Initiated the study, wrote the code and ran the simulations; H.S., S.B., Y.I.W and E.V.K. contributed to the design of the simulations and analyzed the results; H.S. and E.V.K. wrote the manuscript that was edited and approved by all authors.

## Acknowledgements

The authors thank Koonin Group members for helpful discussions. The authors’ research is supported by the Intramural Research Program of the National Institutes of Health of the USA (National Library of Medicine).

