## Supplementary figures and tables for "In silico evolution of globular protein folds from random sequences"

### **Supplementary information**

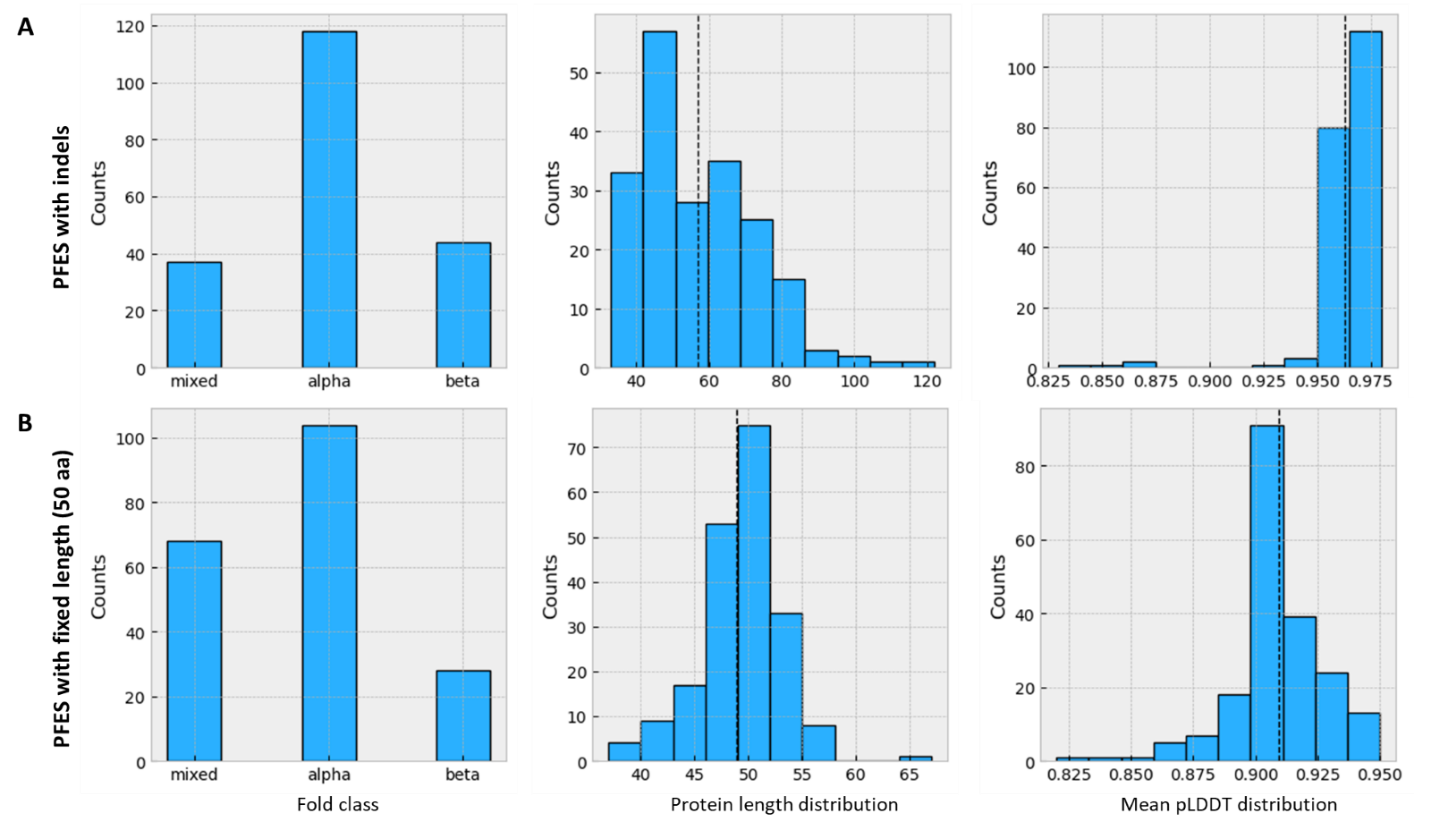

**Figure S1.** Protein fold class, length, mean pLDDT distributions in PFES. Simulations with indels include full or partial duplication and insertions of different lengths and were started from random sequences containing 24 residues. Simulations with constrained protein length were started with random sequences containing 50 residues and only single amino acid indels and substitutions were allowed.

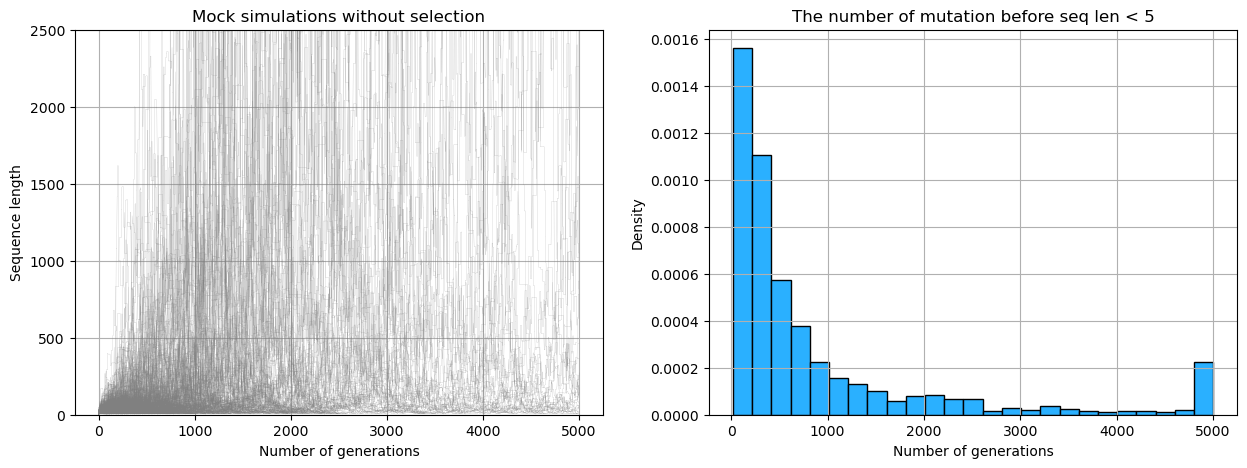

**Figure S2.** PFES without selection demonstrating changes in the protein length driven solely by the rates of indels presented in Table S2. **(A)** Simulations showing the length of sequences in the absence of any selective pressure, n=1000, starting sequence length = 24. **(B)** Distribution of the number of mutations before reaching the sequences sequence length of < 6 residues.

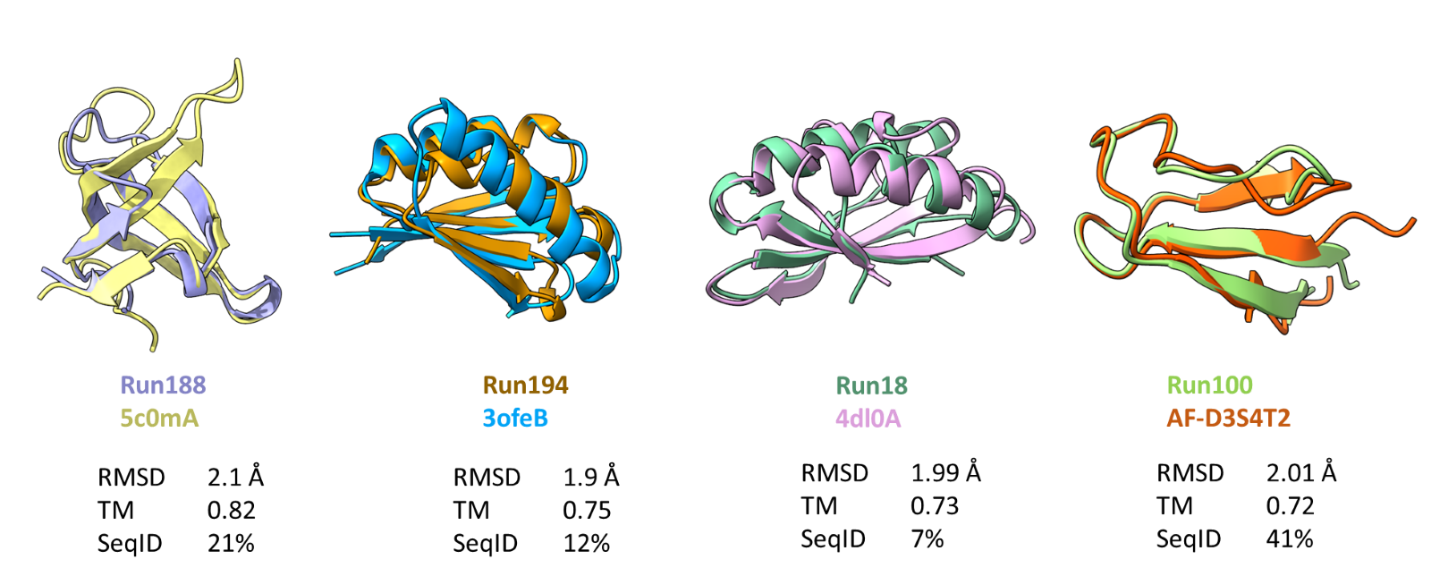

**Figure S3.** Proteins from simulated evolution structurally similar to real proteins from PDB and AFDB databases.

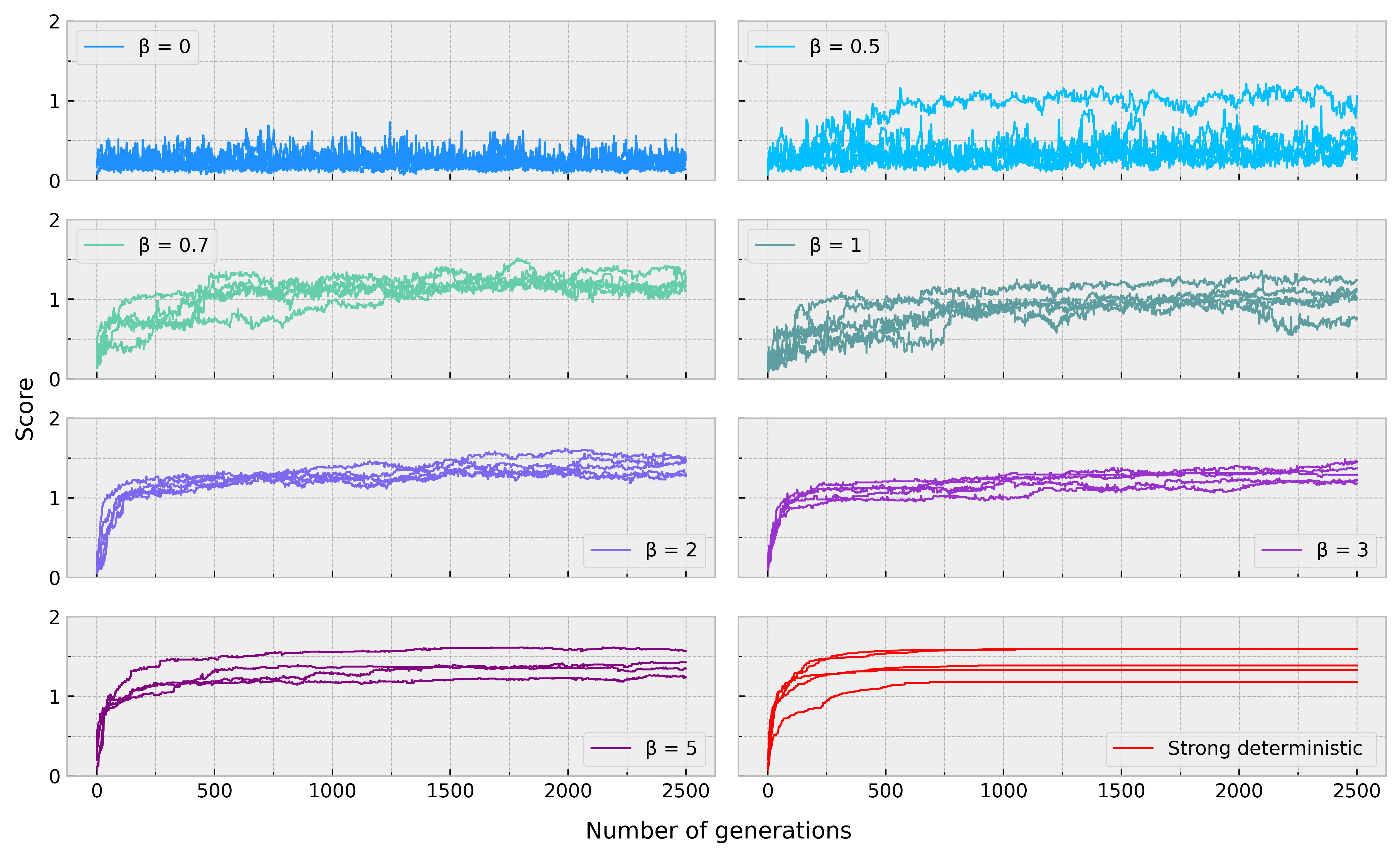

**Figure S4.** Simulation with stochastic selection of a 50 amino acid random peptide with different β.

**Table S1.** Foldseek search results

| **query** | **target** | **prob** | **fident** | **qcov** | **alnlen** | **evalue** | **db** | **nhits** |
| --- | --- | --- | --- | --- | --- | --- | --- | --- |
| run105 | 3vu4_B | 1.00 | 0.08 | 0.90 | 73 | 0.00033 | pdb | 2640 |
| run48 | 7vfk_B | 1.00 | 0.12 | 0.96 | 92 | 0.00180 | pdb | 2023 |
| run69 | 7vfm_C | 1.00 | 0.15 | 0.92 | 73 | 0.00190 | pdb | 2010 |
| run99 | 4x7r_A | 1.00 | 0.19 | 0.82 | 52 | 0.00254 | pdb | 1912 |
| run124 | 2q4m_A | 1.00 | 0.11 | 0.98 | 143 | 0.00350 | pdb | 675 |
| run18 | 4dl0_C | 1.00 | 0.13 | 0.99 | 71 | 0.00748 | pdb | 1071 |
| run93 | 7vfk_B | 1.00 | 0.10 | 0.94 | 68 | 0.00834 | pdb | 1995 |
| run188 | 5c0m_A | 1.00 | 0.18 | 0.98 | 57 | 0.00996 | pdb | 2001 |
| run22 | 5lhz_C | 1.00 | 0.17 | 1.00 | 52 | 0.09474 | pdb | 818 |
| run100 | 6htj_B | 1.00 | 0.19 | 0.97 | 37 | 0.25540 | pdb | 1699 |
| run130 | 6hhm_A | 0.99 | 0.04 | 0.82 | 45 | 0.61880 | pdb | 464 |
| run87 | 5twb_A | 0.99 | 0.09 | 0.84 | 32 | 0.45460 | pdb | 653 |
| run185 | 5twb_A | 0.99 | 0.12 | 0.92 | 34 | 0.46980 | pdb | 776 |
| run16 | 6y2s_A | 0.99 | 0.13 | 0.83 | 45 | 0.53680 | pdb | 580 |
| run77 | 7ywp_A | 0.99 | 0.11 | 0.94 | 84 | 0.04018 | pdb | 417 |
| run133 | 6czi_B | 0.99 | 0.16 | 0.96 | 100 | 0.04278 | pdb | 111 |
| run141 | 3bk5_A | 0.98 | 0.10 | 0.96 | 48 | 0.94090 | pdb | 519 |
| run50 | 3u8v_B | 0.98 | 0.11 | 0.87 | 62 | 0.62310 | pdb | 132 |
| run80 | 2mm3_A | 0.98 | 0.13 | 0.94 | 46 | 0.67640 | pdb | 419 |
| run31 | 5uw7_A | 0.98 | 0.09 | 0.98 | 98 | 0.04906 | pdb | 304 |
| run156 | 6ttu_T | 0.98 | 0.14 | 0.97 | 77 | 0.30910 | pdb | 266 |
| run5 | 2ecf_A | 0.97 | 0.19 | 0.98 | 48 | 0.45540 | pdb | 437 |
| run38 | 7l9e_E | 0.96 | 0.14 | 0.95 | 59 | 0.49280 | pdb | 495 |
| run194 | AF-A0A4Q2RST3 | 1.00 | 0.22 | 1.00 | 69 | 0.00093 | afdb | 651 |
| run41 | AF-A0A143G9F3 | 1.00 | 0.21 | 0.86 | 61 | 0.00479 | afdb | 376 |
| run193 | AF-A0A814L140 | 1.00 | 0.15 | 1.00 | 72 | 0.02876 | afdb | 389 |
| run96 | AF-A0A2D3PMG7 | 1.00 | 0.08 | 0.95 | 89 | 0.03642 | afdb | 131 |
| run199 | AF-A0A7W0LT51 | 1.00 | 0.29 | 0.89 | 42 | 0.04865 | afdb | 816 |
| run146 | AF-A0A7S4L576 | 1.00 | 0.21 | 0.95 | 57 | 0.05781 | afdb | 39 |
| run95 | AF-A0A7C7HK81 | 1.00 | 0.20 | 0.88 | 50 | 0.07864 | afdb | 581 |
| run165 | AF-A0A2D8AN05 | 1.00 | 0.28 | 0.93 | 39 | 0.14870 | afdb | 174 |
| run197 | AF-A0A0F9JEQ6 | 1.00 | 0.16 | 1.00 | 56 | 0.17330 | afdb | 90 |
| run90 | AF-A0A7W7KAE5 | 1.00 | 0.21 | 0.88 | 43 | 0.17680 | afdb | 96 |
| run176 | AF-W7Z7K7 | 1.00 | 0.16 | 0.97 | 38 | 0.17950 | afdb | 499 |
| run180 | AF-A0A0C2W3X4 | 1.00 | 0.05 | 0.92 | 43 | 0.19380 | afdb | 609 |
| run187 | AF-A0A7Y2Y380 | 1.00 | 0.11 | 0.98 | 45 | 0.23340 | afdb | 951 |
| run139 | AF-A0A5E4I478 | 1.00 | 0.19 | 0.96 | 43 | 0.26220 | afdb | 1040 |
| run106 | AF-A0A1Q2CMH5 | 1.00 | 0.24 | 0.96 | 42 | 0.31340 | afdb | 1174 |
| run170 | AF-A0A3M1T0E8 | 1.00 | 0.12 | 1.00 | 66 | 0.31390 | afdb | 268 |
| run102 | AF-A0A495M102 | 1.00 | 0.20 | 0.95 | 35 | 0.35360 | afdb | 31 |
| run92 | AF-A0A7Z9JEL9 | 1.00 | 0.15 | 0.95 | 60 | 0.55210 | afdb | 77 |
| run178 | AF-A0A3D8T9U8 | 1.00 | 0.21 | 0.81 | 43 | 0.61670 | afdb | 74 |
| run111 | AF-A0A3B6THA5 | 1.00 | 0.16 | 0.96 | 45 | 0.50580 | afdb | 21 |
| run171 | AF-A0A7X7Q3P4 | 1.00 | 0.18 | 0.85 | 33 | 0.52200 | afdb | 125 |
| run3 | AF-A0A7X1E4C3 | 1.00 | 0.18 | 0.98 | 87 | 0.02829 | afdb | 9 |
| run115 | AF-A0A5C7PGG2 | 1.00 | 0.12 | 0.89 | 33 | 1.02200 | afdb | 9 |
| run107 | AF-X0SW97 | 1.00 | 0.21 | 0.83 | 47 | 0.89180 | afdb | 59 |
| run186 | AF-A0A3S9TJV0 | 1.00 | 0.10 | 0.98 | 83 | 1.00600 | afdb | 7 |
| run42 | AF-A0A842B3M2 | 1.00 | 0.11 | 1.00 | 45 | 1.20700 | afdb | 3 |
| run29 | AF-A0A0E9NQ65 | 0.99 | 0.21 | 0.90 | 103 | 0.00134 | afdb | 1 |
| run196 | AF-A0A242WAS5 | 0.99 | 0.18 | 0.93 | 57 | 0.23890 | afdb | 16 |
| run140 | AF-A0A414NWH6 | 0.99 | 0.28 | 0.97 | 36 | 2.30300 | afdb | 23 |
| run151 | AF-A0A521K8B3 | 0.99 | 0.33 | 0.91 | 40 | 0.54510 | afdb | 47 |
| run131 | AF-A0A2R6RQF2 | 0.99 | 0.28 | 0.93 | 39 | 1.18800 | afdb | 6 |
| run108 | AF-A0A7C4TWM1 | 0.99 | 0.21 | 0.92 | 56 | 1.62600 | afdb | 2 |
| run190 | AF-A0A1Q4GU02 | 0.99 | 0.15 | 0.83 | 41 | 1.07700 | afdb | 36 |
| run198 | AF-F4P5B8 | 0.99 | 0.19 | 0.96 | 53 | 0.66070 | afdb | 21 |
| run36 | AF-A0A3P6B3K0 | 0.99 | 0.21 | 0.88 | 43 | 2.00800 | afdb | 15 |
| run53 | AF-T1XNQ2 | 0.99 | 0.12 | 0.96 | 82 | 2.01300 | afdb | 3 |
| run25 | AF-A0A849ZRA1 | 0.98 | 0.14 | 0.90 | 43 | 3.90900 | afdb | 1 |
| run129 | AF-I1IM41 | 0.98 | 0.19 | 0.91 | 42 | 4.24100 | afdb | 1 |
| run157 | AF-A0A517TSJ5 | 0.98 | 0.17 | 0.91 | 78 | 1.59100 | afdb | 2 |
| run66 | AF-S1N7D9 | 0.98 | 0.17 | 0.96 | 53 | 4.10100 | afdb | 10 |
| run83 | AF-A0A537X4E9 | 0.98 | 0.14 | 1.00 | 88 | 0.28350 | afdb | 5 |
| run47 | AF-A0A316YHF2 | 0.98 | 0.14 | 0.96 | 90 | 0.37140 | afdb | 7 |
| run128 | AF-A0A1Q6UZD7 | 0.98 | 0.17 | 0.84 | 36 | 1.77100 | afdb | 13 |
| run159 | AF-A0A535GJU4 | 0.97 | 0.08 | 0.93 | 52 | 3.79500 | afdb | 5 |
| run168 | AF-M1VDQ2 | 0.97 | 0.21 | 0.88 | 53 | 4.11700 | afdb | 3 |
| run110 | AF-A0A239QJR5 | 0.97 | 0.21 | 0.83 | 34 | 4.81300 | afdb | 1 |
| run64 | AF-A0A429CVJ2 | 0.96 | 0.25 | 0.95 | 56 | 2.36200 | afdb | 1 |
| run26 | AF-A0A7K4DWC2 | 0.96 | 0.07 | 0.89 | 68 | 3.39600 | afdb | 1 |
| run86 | AF-A0A7J0EWZ7 | 0.96 | 0.17 | 0.98 | 41 | 0.86610 | afdb | 4 |
| run109 | AF-A0A418PM05 | 0.96 | 0.25 | 0.82 | 28 | 4.14700 | afdb | 2 |
| run32 | MGYP000472187349 | 0.99 | 0.15 | 0.92 | 62 | 0.17890 | esm | 1 |
| run72 | MGYP001476248561 | 0.99 | 0.16 | 0.92 | 55 | 1.59300 | esm | 4 |
| run88 | MGYP003324074694 | 0.98 | 0.20 | 0.95 | 41 | 0.24300 | esm | 2 |
| run94 | MGYP001385367156 | 0.98 | 0.14 | 0.95 | 78 | 0.70690 | esm | 1 |
| run52 | MGYP003295344537 | 0.98 | 0.18 | 0.91 | 40 | 1.33500 | esm | 1 |
| run104 | MGYP003590916796 | 0.98 | 0.16 | 0.98 | 62 | 0.47420 | esm | 1 |
| run154 | MGYP003706959411 | 0.96 | 0.16 | 0.97 | 69 | 1.54600 | esm | 1 |
| run79 | MGYP003353214145 | 0.96 | 0.13 | 0.87 | 32 | 1.75000 | esm | 1 |

rep target – a representative target for this query; prob – foldseek probability; fident – fraction of identical matches; qcov – query coverage, alnlen – alignment length; db – database where the representative hit is coming from; nhits – total number of hits

**Table S2. All Foldseek hits with probablity > 0.95 and query coverage of 80%.**

**Tables S3.**

| Amino acid | AA | Flat rate | Codon rates | Uniprot rate |
| --- | --- | --- | --- | --- |
| Alanine | A | 1 | 1.311 | 1.652 |
| Cysteine | C | 1 | 0.656 | 0.278 |
| Aspartate | D | 1 | 0.656 | 1.092 |
| Glutamate | E | 1 | 0.656 | 1.344 |
| Phenylalanine | F | 1 | 0.656 | 0.774 |
| Glycine | G | 1 | 1.311 | 1.414 |
| Histidine | H | 1 | 0.656 | 0.456 |
| Isoleucine | I | 1 | 0.984 | 1.182 |
| Lysin | K | 1 | 0.656 | 1.16 |
| Leucine | L | 1 | 1.967 | 1.93 |
| Methionine | M | 1 | 0.328 | 0.482 |
| Asparagine | N | 1 | 0.656 | 0.812 |
| Proline | P | 1 | 1.311 | 0.95 |
| Glutamine | Q | 1 | 0.656 | 0.786 |
| Arginine | R | 1 | 1.967 | 1.106 |
| Serine | S | 1 | 1.967 | 1.33 |
| Threonine | T | 1 | 1.311 | 1.072 |
| Valin | V | 1 | 1.311 | 1.372 |
| Tryptophan | W | 1 | 0.328 | 0.22 |
| Tyrosine | Y | 1 | 0.656 | 0.584 |
| Single residue insertion | + | 1 | 1 | 1 |
| Single residue deletion | - | 1 | 1 | 1 |
| Partial duplication | * | 0.4 | 0.4 | 0.4 |
| Random insertion | / | 0.4 | 0.4 | 0.4 |
| Partial deletion | % | 0.9 | 0.9 | 0.9 |
| Circular permutation | p | 0.1 | 0.1 | 0.1 |
| Full duplication | d | 0.05 | 0.05 | 0.05 |
